# Polyphosphate initiates tau aggregation through intra- and intermolecular scaffolding

**DOI:** 10.1101/588509

**Authors:** S. P. Wickramasinghe, J. Lempart, H. E. Merens, J. Murphy, U. Jakob, E. Rhoades

## Abstract

The aggregation and deposition of tau is a hallmark of a class of neurodegenerative diseases called tauopathies. Despite intensive study, cellular and molecular factors that trigger tau aggregation are not well understood. Here we provide evidence for two mechanisms relevant to the initiation of tau aggregation in the presence of cytoplasmic polyphosphates (polyP): changes in the conformational ensemble of monomer tau and noncovalent cross-linking of multiple tau monomers. We identified conformational changes throughout full-length tau, most notably diminishment of long-range interactions between the termini coupled with compaction of the microtubule binding and proline rich regions. We found that while the proline rich and microtubule binding regions both contain polyP binding sites, the proline rich region is a requisite for compaction of the microtubule binding region upon binding. Additionally, both the magnitude of the conformational change and the aggregation of tau are dependent on the chain length of the polyP polymer. Longer polyP chains are more effective at intermolecular, noncovalent cross-linking of tau. These observations provide an understanding of the initial steps of tau aggregation through interaction with a physiologically relevant aggregation inducer.

## Introduction

Neurodegenerative tauopathies are a class of heterogeneous dementias and movement disorders characterized by abnormal accumulation of the microtubule-associated protein tau in insoluble fibrillar aggregates (1). Aggregation transforms soluble, unstructured tau monomers into highly insoluble, β-sheet rich paired helical filaments and neurofibrillary tangles. While the increase of these insoluble aggregates and the progression of symptoms of neurodegeneration are linked, the triggers and mechanisms by which aggregation occurs in the brain are not well characterized.

Normally, tau binds to and stabilizes microtubules and plays an important role in the organization of the cytoskeleton of neuronal cells (2). Monomer tau is intrinsically disordered and remains largely disordered even upon binding to microtubules (3, 4). *In vitro*, tau is highly soluble and aggregates slowly; polyanionic molecules, such as the extracellular matrix glycosaminoglycan heparin, arachidonic acid or lipid vesicles are commonly used to enhance the rate of aggregation (5-7). We recently demonstrated a similar activity for linear chains of anionic phosphates (i.e. polyphosphate, polyP), which dramatically enhanced the aggregation rates of several amyloidogenic proteins, including tau (8). PolyPs are present at micromolar concentrations in the cytosol of mammalian neurons (9, 10) where they have the potential to interact with tau under normal physiological conditions.

Tau aggregation is generally described as nucleation-dependent, with the nucleating species thought to be an “aggregation-prone” monomer (11). Recent evidence in support of this model comes from a study which identified monomer tau derived from heparin-induced aggregates as capable of seeding tau aggregation both *in vitro* and in cultured cells (12). A molecular description of the conformational change in monomer tau that predisposes it towards aggregation is of critical importance to developing a complete picture of the molecular determinants of aggregation. However, structural characterization of monomer tau is challenging due to its dynamic, disordered nature. Nuclear magnetic resonance (NMR), which is capable of generating amino acid specific structural information(13), requires high protein concentrations that favor rapid aggregation in the presence of molecular inducers. Work from our lab has established that single molecule Förster resonance energy transfer (FRET) is a powerful approach to characterize conformations of aggregation-prone proteins (14, 15).

In this study, we used single molecule FRET and fluorescence correlation spectroscopy (FCS) to investigate polyP binding to tau. We identified that both the microtubule binding region (MTBR) and the proline rich region (PRR) (Fig. 1) contain binding sites for polyP. Moreover, we found that longer polyP polymers were more effective at accelerating tau aggregation than shorter ones. Based on our results, we propose that polyP can serve both as an intramolecular scaffold, by binding to multiple sites within a single tau molecule, as well as an intermolecular scaffold, by binding to multiple tau molecules simultaneously.

**Figure 1.**
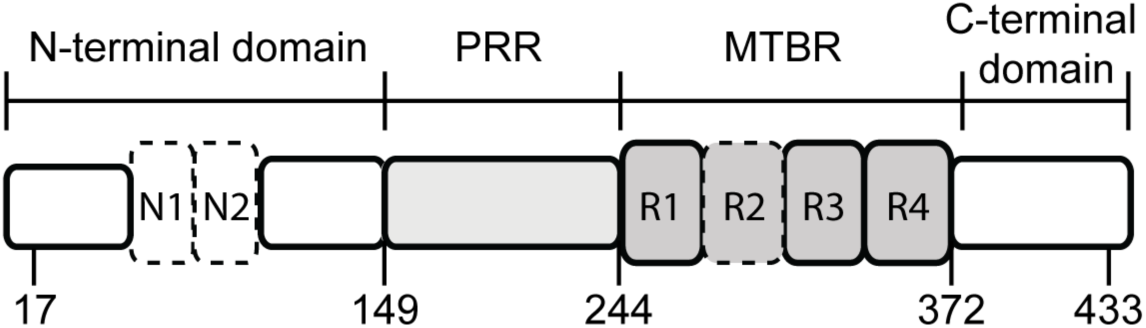
Tau Schematic. The longest isoform of tau, 2N4R, is shown. Regions of interest indicated are the N-terminal domain, the proline rich region (PRR), the microtubule binding region (MTBR) and the C-terminal domain. The residues mutated to cysteine for labeling span these regions of interest and are indicated in the schematic. Alternative splicing of N1, N2 and R2 (marked with dashed lines) result in the six isoforms of tau found in human adults.

## Materials and Methods

### Protein purification and labeling

Tau constructs were purified based on previously published protocols (15, 16). Briefly, all variants were expressed with a cleavable N-terminal His-tag. After elution from a nickel column with 400mM mM imidazole, the imidazole concentration was reduced through several buffer exchange cycles using Amicon concentrators (Millipore) and the protein was incubated with tobacco etch virus (TEV) protease at 4 °C overnight. After a second nickel column to remove the enzyme and the cleaved tag, the protein was further purified by size-exclusion chromatography on a HiLoad 16/600 Superdex 200 Column (GE Life Sciences).

For site-specific labeling with fluorophores, QuickChange mutagenesis was used to change both native cysteines to serines and to introduce new cysteines at desired locations. For single molecule FRET measurements, cysteines were chosen to span domains of interest as described in the manuscript. For FCS measurements, the protein was labeled with a single cysteine introduced near either the N- or C-terminus. For labeling, freshly purified tau was first reduced with 1 mM DTT for 10 minutes and buffer exchanged into a labeling buffer (20 mM Tris pH 7.4, 50 mM NaCl, and 6 M guanidine HCl) using Amicon concentrators (Millipore) (17). For labeling with donor and acceptor fluorophores for single molecule FRET, protein was incubated stirring at room temperature for an hour with donor fluorophore Alexa Fluor 488 maleimide (Invitrogen) at a 2:1 protein:dye ratio. The acceptor fluorophore, Alexa Fluor 594 maleimide (Invitrogen) was then added in 5-fold molar excess and incubated stirring overnight at 4°C. For single labeling for FCS, Alexa Fluor 488 maleimide was added in 5-fold molar excess to protein and incubated overnight stirring at 4 °C. Labeled protein was buffer exchanged into 20 mM Tris (pH 7.4) and 50 mM NaCl and unreacted dye was removed by passing the solution over a HiTrap Desalting Column (GE Life Sciences).

### Aggregation Assays

Fibril formation of tau fragments was monitored using Thioflavin T fluorescence. 25 µM tau and 10 µM Thioflavin T were incubated in 40 mM potassium phosphate, 50 mM KCl pH 7.5 at 37 °C with 1 mM of different chain length of polyP (polyP kindly provided by T. Shiba (Regenetiss, Japan)) or 18 µM heparin from porcine intestinal mucosa (molecular weight 17–20 kDa; Sigma-Aldrich). The polyP concentration is in monomer phosphate units and the heparin concentration was chosen to match the amount of negative charge of the 1 mM polyP. Approximate equivalent charge concentrations of heparin were calculated assuming 1.5 to 2 charges and an average molecular weight of 665 g/mol per disaccharide (18). The samples were agitated for 10 seconds before every reading. The measurements were made in a black 96-well polystyrene microplate with clear bottom (Greiners). Experiments were read in a Synergy HTX MultiMode Microplate Reader (Biotec) with an excitation wavelength of 430 nm and emission detected at 485 nm.

### Single molecule FRET instrument and data analysis

Single molecule FRET measurements were performed on a lab-built instrument based on an inverted Olympus IX-71 Microscope (Olympus) (15, 16). The laser power (488-nm diode-pumped solid-state laser, Spectra-Physics) was adjusted to 25-35 µW before entering the microscope. Fluorescence emission was collected through the objective, and photons were separated by an HQ585LP dichroic in combination with ET525/50M and HQ600LP filters for the donor and acceptor photons, respectively (all filters and dichroics from Chroma). Fluorescence signals were collected by 100 µm diameter aperture fibers (OzOptics) coupled to avalanche photodiodes (Perkin-Elmer). Photon traces were collected in 1 ms time bins for an hour.

All measurements were carried out at a protein concentration of ∼30 pM at 20 °C in 40 mM potassium phosphate, 50 mM KCl buffer pH 7.4 in 8-chambered Nunc coverslips (Thermo-Fisher) passivated with poly(ethylene glycol) poly(L-lysine) (PEG-PLL) to reduce protein and polyP adsorption to the chamber.

To discriminate photon bursts of real events from background noise, a threshold of 30 counts/ms for the sum of the donor and acceptor channels was applied (14). Measurements were made of polyP samples in the absence of tau to determine the background signal (see Eq. 2 below) as well as any spurious contribution to single molecule FRET events. No photon bursts (as defined by the criteria above) were observed for hour-long measurements of 20 µM polyP.

The photon traces were analyzed using MATLAB (Mathworks) based lab-written software. For each event, the energy transfer efficiency (ET_eff_) value was calculated from:

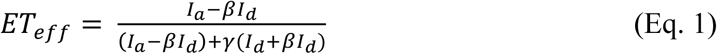

where *I*_*a*_ and *I*_*d*_ are the fluorescence intensities collected in the acceptor and donor channels, respectively. β accounts for fluorescence from the donor fluorophore on the acceptor channel, and is measured each day varying between 0.7 and 0.85 for both the home-built and commercial systems. γ accounts for differences in detection efficiency and quantum yield for acceptor and donor fluorophores (19) and is measured every few months; γ values used on the lab-built system were 1.36, 1.20 and 1.00 (changes in γ occurred after major realignments or other adjustments to the instrument) and were 1.20 and 1.08 on the commercial Picoquant system (value changed after reinstalling the laser). These individual ET_eff_ values were compiled into histograms and fit by multi-peak Gaussian functions to determine properties of the distributions. We compared ET_eff_ histograms calculated with a correction for the background signal (Eq. 2) (14) with those calculated without one (Eq. 1)

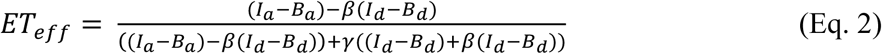

Fitting of the histograms yielded equivalent mean ET_eff_ values (Fig. S1).

Double-labeling of the proteins usually yields a mixture of labeled proteins with donor-only, acceptor-only and donor-acceptor populations. The donor- and acceptor-only labeled are easily separated in analysis. Donor-only labeled protein contributes to the ‘zero-peak’ (ET_eff_ = 0) and acceptor-only labeled proteins does not give rise to a signal as the acceptor is not directly excited (17).

For constructs where the major protein peak had a low ET_eff_ and thus was difficult or impossible to distinguish from the donor-only ‘zero-peak’, direct excitation of the acceptor fluorophore was used to discriminate these two populations (20). Measurements were made in pulsed interleaved excitation FRET (PIE-FRET) mode on a MicroTime 200 inverse time-resolved confocal microscope (Picoquant). Laser power from 485 and 560 nm lasers pulsed at 40 MHz, were adjusted to be ∼30 µW before entering the microscope. Fluorescence emission was collected through the objective and passed through a 100 µm pinhole. Photons were separated by an HQ585LP dichroic in combination with ET525/50M and HQ600LP filters and collected on photodiodes. ET_eff_ and stoichiometry factors were calculated using SymphoTime 64 software and the resultant histograms were fit using Gaussian distributions as described above. Both the lab-built and Microtime 200 instruments were calibrated using 10 base pair, 14 base pair and 18 base pair dsDNA standards.

The saturating concentration of polyP was determined by a titration of polyP_300_ probing the N-terminal domain (tau_C17-C149_) and the PRR and MTBR (tau_C149-C372_) (Fig. S2). The intermediate concentrations showed the rise of double peaks, suggesting heterogeneous populations. The peak positions stops shifting after 20 µM polyP of two different chain lengths (300 and 60 phosphate subunits). These titrations were also repeated with the 3R isoforms.

### FCS instrument and data analysis

On the lab-built instrument, the laser power was adjusted to ∼5 µW as measured prior to entering the microscope. Fluorescence emission was collected through the objective and separated from laser excitation using a Z488RDC long pass dichroic and an HQ600/200M bandpass filter and focused onto a 50 µm diameter optical fiber directly coupled to an avalanche photodiode. A digital correlator (FLEX03LQ-12, Correlator.com) was used to generate the autocorrelation curves. All FCS measurements were carried out at ∼20 nM protein at 20 °C in PEG-PLL coated Nunc coverslips. For each measurement, 25 traces of 10 seconds were averaged to obtain statistical variations and then fit to a diffusion equation using a single-component fit using lab-written scripts in MATLAB (The Mathworks).

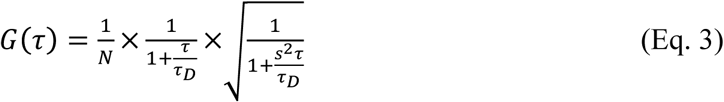

Some measurements were carried out on the MicroTime 200 instrument and analyzed using the SymphoTime 64 software.

Noncovalent cross-linking measurements were carried out using 25 µM unlabeled 4R, 20 nM single labeled 4R and 1 mM polyP or 18 µM heparin at 20 °C. Triplicate measurements with 25 curves of 10 seconds were recorded immediately after mixing. The cross-linking FCS measurements were carried out at a lower temperature (20 °C) than the aggregation assays (37 °C). Aggregation occurs more slowly at lower temperatures such that it is unlikely that large fibrillar aggregates are formed during the duration of the FCS measurements.

### Competition Assays

Tubulin was purified from bovine brain tissue using repeated cycles of polymerization and depolymerization in the presence of a high-molarity PIPES buffer (21). Before use, tubulin was clarified by centrifugation at 100,000g and buffer-exchanged into phosphate buffer.

Competitive binding of tubulin and polyP to tau was measured by FCS. ∼20 nM P2-4R was incubated with 5 µM tubulin for 5 minutes at 20 °C. PolyP_300_ was added to the tau-tubulin sample at concentrations ranging from 1 µM to 100 µM, and incubated for 5 minutes before measurement. For each measurement, 25 curves of 10 seconds were recorded and analyzed with a single-component fit (Eq. 3 above) using lab-written scripts in MATLAB. Measurements were repeated in triplicate.

### Statistical Analysis

Data are presented as mean ± s.e.m. Statistical significance was calculated using unpaired two-tailed tests with P<0.05 considered significant with a minimum of n=3 for FRET, FCS and aggregation assays.

## Results

Functionally, tau can be divided into four major domains (Fig. 1). The function of the N-terminal domain is the least well-understood, although it has been proposed to regulate binding to microtubules (22) as well as to interact with neuronal membranes (23). The adjacent MTBR binds both microtubules and soluble tubulin (24, 25) and forms the core of paired helical filaments. Binding of tau to microtubules is enhanced by the MTBR-flanking PRR and the C-terminal domain (26, 27). Alternate splicing of the second repeat (R2) in the MTBR gives rise to isoforms with either three (3R) or four (4R) repeat regions and either zero (0N), one (1N) or two (2N) N-terminal inserts (Fig. 1) (28). The 3R and 4R isoforms have different binding affinities (29) and assembly activities (30) for microtubules. One of the two hexapeptide sequences that play a crucial role in tau aggregation is located in R2. Since this region is absent in 3R isoforms, 4R isoforms are generally more aggregation-prone (31, 32).

Changes to the conformations of each of the domains caused by polyP binding to tau were probed with single molecule FRET. For site-specific labeling of tau in these measurements, cysteine residues were introduced at desired labeling positions in the 2N4R and 2N3R isoforms of tau (Fig. 1; details in *Materials and Methods*). The labeling positions were chosen to span either the entire protein (C17-C433) or specific regions (all residue numbering throughout is based on the 2N4R isoform): the N-terminal domain (C17-C149); PRR and MTBR (C149-C372); or the C-terminal domain (C372-C433). The donor and acceptor fluorophores were Alexa Fluor 488 maleimide and Alexa Fluor 594 maleimide, respectively.

FRET efficiencies (ET_eff_) from individual photon bursts were calculated as a ratio of the intensity of the acceptor over the sum of the intensities of the donor and acceptor (see *Materials and Methods* for details and correction factors). Higher ET_eff_ values correspond to shorter distances between the donor and acceptor dyes, whereas lower ET_eff_ values reflect longer distances. ET_eff_ values were plotted as histograms and fit with a sum of Gaussian distributions to determine the peak ET_eff_ positions (Fig. 2). As an intrinsically disordered protein, tau populates an ensemble of conformations. For simplicity and convenience in comparing FRET measurements of different tau constructs or under different experimental conditions, we use the mean ET_eff_ obtained from fitting to reflect the average of tau’s conformational ensemble for a given measurement. Measurements were made in the absence (Fig. 2, dashed lines) or presence of polyP_300_ (average of 300 phosphate units per polymer) (Fig. 2, solid lines); polyP concentrations are given in monomer P_i_ units.

**Figure 2.**
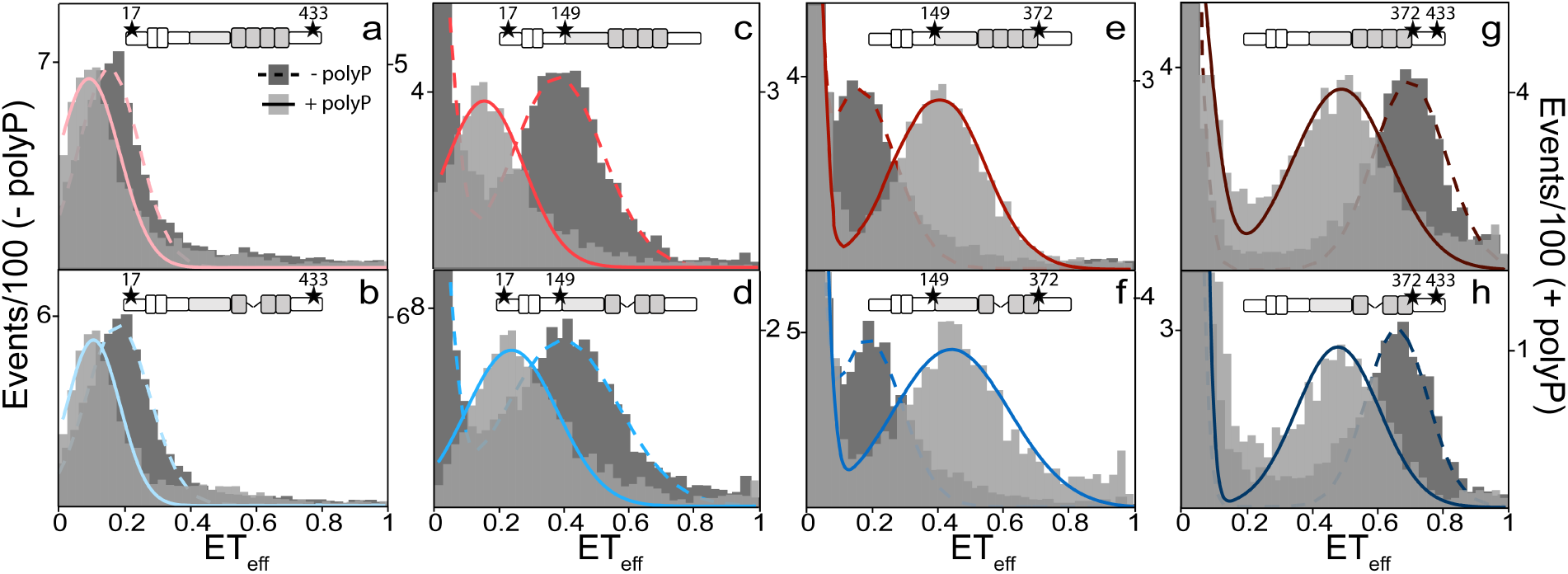
PolyP disrupts long-range interactions and compacts the PRR and MTBR of tau. Histograms from smFRET measurements of 2N4R (upper panels) and 2N3R (lower panels) tau in the absence (dark gray and dashed line; left axis) and presence (light gray and solid line; right axis) of 20 µM polyP_300_. Labeling positions are chosen to probe the entire protein (C17-C433: a, b); the N-terminal domain (C17-C149: c, d); the PRR and MTBR (C149-C372: e, f); and the C-terminal domain (C372-C433: g, h). The numbers are the labeling positions by residue number based on 2N4R. Tau schematics above each histogram represent the isoform and labeling positions. At least three separate measurements of each condition/construct were made. The histograms shown are representative. Statistical analysis of repeat measurements is given in Table S1.

### PolyP disrupts long-range intramolecular interactions in tau

In solution, tau is a compact protein despite its length and intrinsic disorder, with the N- and C-termini in relatively close proximity to the MTBR (15). 2N3R is 31 residues shorter than 2N4R due to the absence of R2; thus, differences in the measured ET_eff_ between the two isoforms for constructs that span this region, i.e., tau_C149-C372_ (MTBR and PRR) and tau_C17-C433_ (entire protein) are expected and observed (Fig. 2; Table S1). Interestingly, the construct probing the C-terminal domain tau_C372-C433_, which is identical in both isoforms, shows a lower ET_eff_ for 2N3R than 2N4R, reflecting a more extended conformational ensemble for 2N3R. One possible explanation is that attractive electrostatic interactions between the C-terminal domain and the MTBR are weakened by the absence of R2.

The addition of polyP_300_ resulted in an increase in the average end-to-end distance of tau as reflected by the decrease in ET_eff_ for both 2N4R (Fig. 2a) and 2N3R (Fig. 2b) tau_C17-C433_ (Table S1). These results suggested that polyP disrupts some of the long-range electrostatic interactions that are responsible for tau’s relatively compact conformational ensemble in solution. We also noted a similar polyP–mediated shift to lower ET_eff_ values, i.e., larger distances, for both isoforms when the N- and C-terminal domains were probed independently, using tau_C17-C149_ or tau_C372-C433_, respectively (Fig. 2c, d and Fig. 2g, h; Table S1). In contrast, tau_C149-C372,_ which probes the region encompassing the PRR and MTBR, showed a relatively large shift towards higher ET_eff_ values for both isoforms, reflecting significant compaction of this region (Fig 2e, f; Table S1). Increasing the ionic strength of the buffer solution to 500 mM KCl eliminated the observed changes in the ET_eff_ histograms for 2N4R tau_C17-C149_ or tau_C149-C372_ (Fig. S3), supporting the idea that electrostatic interactions are important for polyP binding to tau. These results are similar to our previous results on tau-heparin interactions (15), suggesting that there are conserved features in the aggregation-prone conformational ensemble of full-length monomer tau in the presence of polyanionic aggregation inducers.

### PolyP binds to the PRR and MTBR

The MTBR carries an effective net charge of +10.2 and +7.1 at pH 7.4 in the 2N4R and 2N3R isoforms, respectively, while the PRR has a net charge of +13.8. Thus, our expectation was that the negatively charged polyP binds to these domains and that changes in the conformational ensembles in other regions of the protein may result from screening of the positive MTBR and PRR by the anionic polyP. For example, binding of polyP to the MTBR or PRR may disrupt the long-range interactions with the negatively charged N-terminal domain, thereby altering its conformational ensemble even if polyP does not bind directly to the N-terminal domain. To identify the regions of tau involved in polyP binding, we made six tau fragments (Fig. 3): 4R and 3R, which correspond to the MTBR (residues 244-372) of 2N4R and 2N3R, respectively; P1P2, which is the entire PRR (residues 148-245); P2-4R and P2-3R, which contain 4R and 3R as well the second half of the PRR (residues 198-372); and NT, which corresponds to the N-terminal domain (residues 1-152). The fragments and full-length tau constructs were labeled with a single fluorophore and fluorescence correlation spectroscopy (FCS) was used to measure their diffusion times in the absence or presence of polyP_14_ or polyP_300_. A change in mass or hydrodynamic radius of tau upon binding polyP is expected to result in a change in tau’s diffusion time (Fig. S4; details in Materials and Methods). All constructs except for NT (which showed no change) exhibited an increase in diffusion time ranging from 11 to 24% in the presence of polyP_300_ relative to the diffusion time of the construct in the absence of polyP_300_, indicative of polyP binding (Fig. 3; Table S2). The two fragments lacking R2 in their MTBRs (i.e., P2-3R and 3R), however, showed reproducibly less of an increase in diffusion time upon binding polyP than their 4R counterparts (i.e., P2-4R and 4R), suggesting that presence of R2 enhances the interactions with polyP (Fig. 3). Moreover, addition of the shorter polyP_14_ caused similar increases in the diffusion times although of smaller magnitudes than observed for polyP_300_ (Fig. 3). Based on these results, we concluded that polyP binds to both the PRR and MTBR. In addition, the observed lack of polyP binding to the NT indicated that the large conformational changes that we observed when we probed the N-terminal domain in the context of the full-length protein, tau_C17-C149_ (Fig. 2c, d) is not due to direct binding of polyP to this domain but caused by an overall reconfiguration of tau’s conformational ensemble upon binding of polyP to the PRR and MTBR.

**Figure 3.**
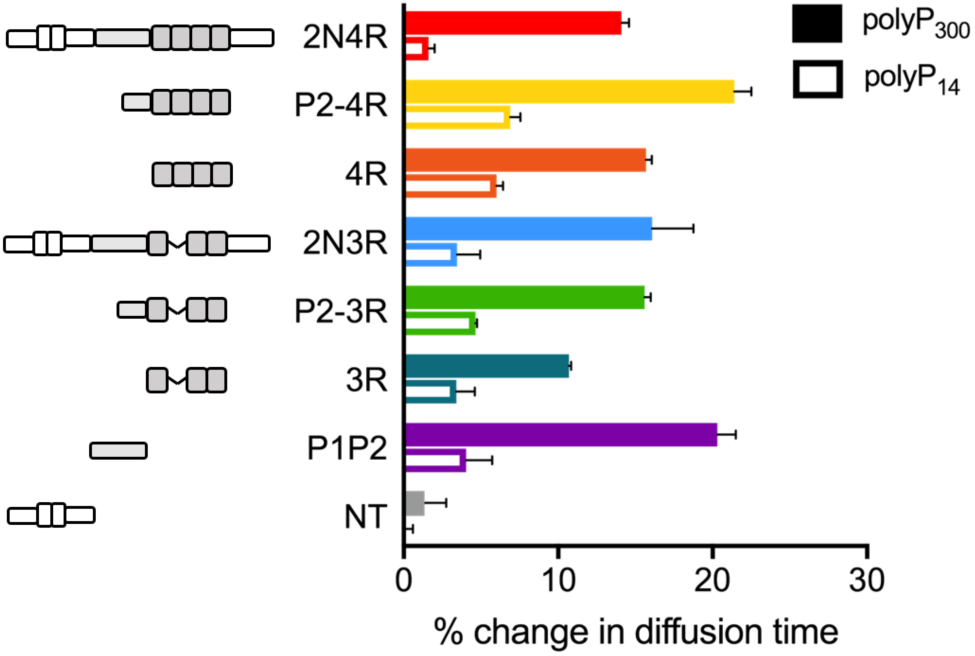
PolyP binds to the PRR and MTBR. Change in diffusion times of single labeled tau constructs with the addition of 20 µM polyP_300_ (solid bars) or polyP_14_ (open bars) as measured by FCS. The change is shown as the increase relative to each construct in the absence of polyP. The results are the average of three separate measurements, and error bars show propagated s.e.m. The changes in diffusion time between 4R and 3R, and P2-3R and 3R are significant (p=0.016 and p=0.05 respectively). Statistical analysis of repeat measurements is given in Table S2.

To understand the relationship between polyP’s binding to the PRR and MTBR and the conformational changes that we observed by smFRET (Fig. 2), we conducted smFRET measurements focusing on either the MTBR (Fig. 4a-c for 4R; Fig. 4d-f for 3R) or the PRR (Fig. 4g-i) in the context of the full-length constructs or select fragments. In the absence of polyP, the fragments generally showed higher mean ET_eff_ values than the full-length proteins, indicating that the flanking regions of the protein impact the conformational ensembles sampled by the various domains even in solution (Fig. 4, dashed lines; Table S3). Full-length tau_C244-C372_ constructs exhibited a large shift to higher ET_eff_ upon binding polyP, reflecting a polyP-induced compaction of the MTBR (Fig. 4a, d; Table S3). We observed similar results for P2-4R_C244-C372_ and P2-3R_C244-C372_ constructs (Fig. 4b, e; Table S3), although the magnitudes of the shifts were not as great as those seen in the full-length proteins. Interestingly, polyP did not cause any significant shifts in the ET_eff_ values in either of the isolated MTBR fragments, 4R_C244-C372_ or 3R_C244-C372_ (Fig. 4c, f; Fig. S5 and Table S3) despite clear evidence of binding to this region as measured by FCS (Fig. 3). However, comparison of the ET_eff_ histograms revealed that binding of polyP to both the full-length tau or the shorter P2-4R_C244-C372_ and P2-3R_C244-C372_ constructs shifts their peak positions close to those of the isolated MTBR fragments, 4R_C244-C372_ or 3R_C244-C372_. These results suggest that the MTBR fragments in isolation populate a compact monomer conformational ensemble that is only achieved in the longer tau constructs through binding of polyP. Since isolated MTBR fragments aggregate more readily than full-length tau (31), our results suggest that compaction of this domain may play a role, reflecting one mechanism by which polyP accelerates tau aggregation.

**Figure 4.**
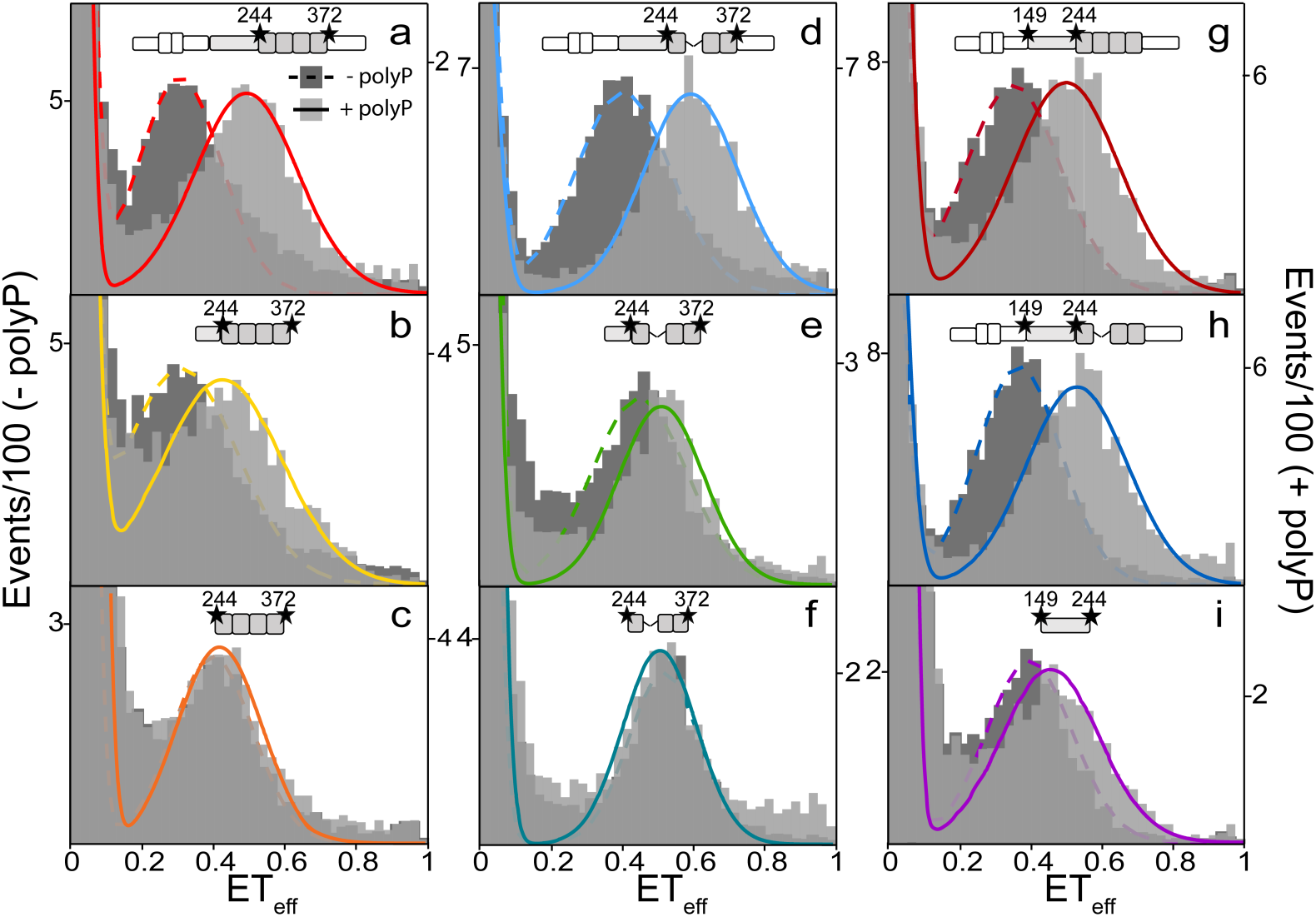
PolyP binding induces conformational changes in the MTBR when the PRR is present. Histograms from smFRET measurements probing the MTBR and PRR in the absence (dark gray and dashed line; left axis) or presence (light gray and solid line; right axis) of 20 µM polyP_300_. Histograms for probes of the MTBR (C244-C372) in 2N4R (a), P2-4R (b), 4R (c), 2N3R (d), P2-3R (e) and 3R (f). Histograms for probes of the PRR (C149-C244) for 2N4R (g), 2N3R (h) and P1P2 (i). Tau schematics above each histogram represent the isoform and labeling positions. At least three separate measurements of each condition/construct were made. The histograms shown are representative. Statistical analysis of repeat measurements is given in Table S3.

SmFRET measurements of the other tau fragments were consistent with FCS measurements that showed binding to P1P2 but not NT. PolyP binding to the isolated PRR, P1P2_C149-C244_ caused a shift to higher ET_eff_ (Fig. 4i; Table S3), although not as great as seen in full-length tau_C149-C244_(Fig. 4g, h; Table S3). No shift in ET_eff_ was detected in the isolated N-terminal fragment, NT_C17-_ C149 (Fig. S5; Table S3).

### PolyP enhances tau aggregation through intermolecular cross-linking

*In vivo*, polyP exists as a broad range of polymer lengths (10) and our prior work with polyP revealed that longer chains are generally more effective at accelerating aggregation of amyloidogenic proteins than shorter ones (8). To examine whether a relationship exists between polyP chain length, changes to the conformational ensemble of the monomer protein and aggregation propensity, we conducted smFRET measurements on select full-length 2N4R constructs, tau_C17-C149_ (Fig. 5a) and tau_C149-C372_ (Fig. 5b) in the presence of saturating concentrations of polyP_14_, polyP_60_, polyP_130_ or polyP_300_ (Table S4). In the presence of polyP_14_, tau_C149-C372_ underwent a very small increase in ET_eff_ while the tau_C17-C149_ conformation appeared to be unchanged (Fig. 5; Table S4). This was despite the fact that polyP_14_ binds to both the full-length protein as well as to the isolated fragments (Fig. 3). All other polyP chain lengths caused large shifts in the mean ET_eff_ for both tau_C17-C149_ and tau_C149-C372_ to the previously observed lower and higher values, respectively (Fig. 5; Table S4). This suggests that for longer polyP polymers, a single chain may be able to bind to multiple binding sites on tau to allow for intra-molecular scaffolding resulting in a change in its conformational ensemble. When the chain is not long enough (i.e. polyP_14_) this effect is not observed.

**Figure 5.**
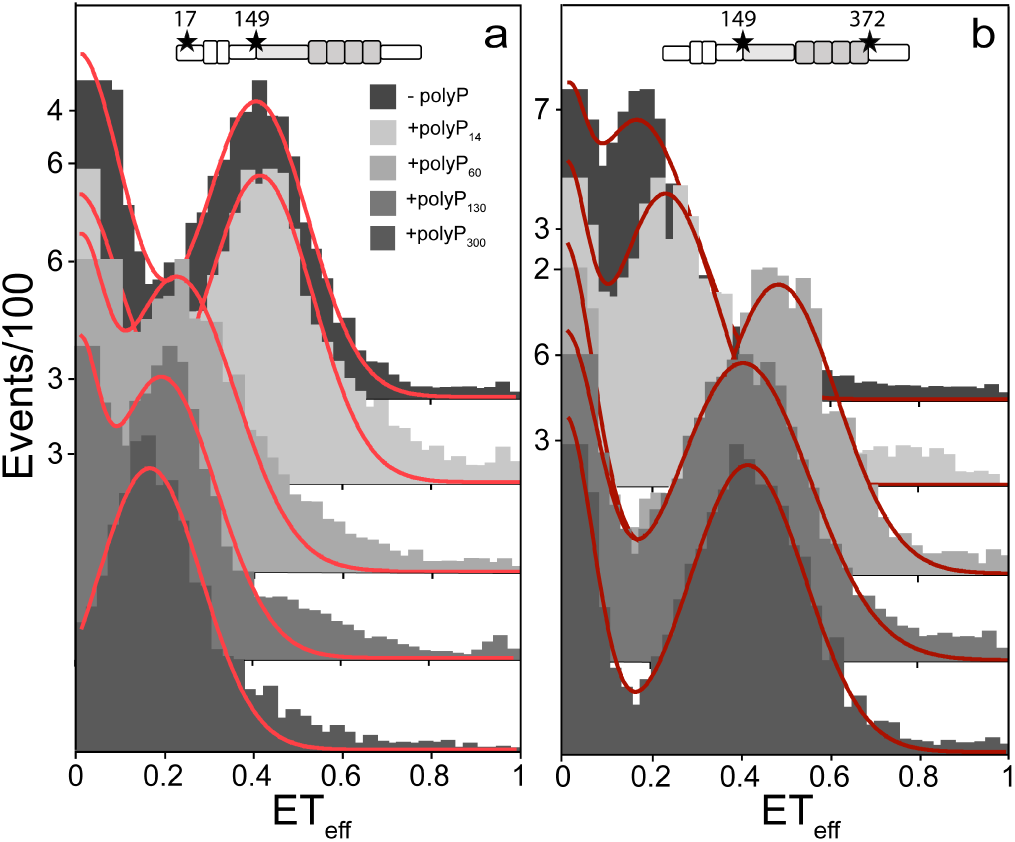
Tau conformational changes are dependent upon polyP chain length. Histograms from smFRET measurements probing two different regions of 2N4R, the N-terminal domain (C17-C149) (a) and the PRR and MTBR (C149-C372) (b) in the absence or presence of 20 µM polyP of different chain lengths. At least three separate measurements of each condition/construct were made. The histograms shown are representative. Statistical analysis of repeat measurements is given in Table S4.

Ensemble aggregation experiments were carried out for tau fragments in the presence of various chain lengths of polyP. Aggregation was monitored by an increase in Thioflavin T fluorescence and quantified by the T_1/2_, the time to reach half the signal plateau (Fig. 6; Fig. S6 and Table S5; details in Materials and Methods). While an inverse relationship between the T_1/2_ and the length of polyP was observed for both 3R and 4R, all polyP chain lengths were more effective at inducing aggregation than heparin (Fig. 6). To illustrate, while the T_1/2_ of 4R was over 15 hours in the presence of heparin, it was less than 6 hours in the presence of polyP_14_ and less than 10 minutes in the presence of polyP_300_. Consistent with previous findings that the 4R-fragment of tau is generally more aggregation-prone than the 3R-fragment (33) (e.g., T_1/2_ of 15 hours *versus* 60 hours in the presence of heparin), the influence of all polyP chains was accordingly less pronounced for the 3R-fragment. Moreover, while polyP was effective in accelerating the aggregation of the two 4R constructs (R4 and P2-R4), it was less effective at inducing aggregation of 3R as compared to P2-3R (Fig. 6; Table S5). These data, along with the binding data shown in Fig. 3, support the importance of polyP binding to both R2 and the PRR for effective acceleration of aggregation; constructs which contain either P2 (i.e., P2-3R), R2 (i.e, 4R) or both (i.e., P2-4R) aggregate significantly faster in the presence of polyP than 3R, which lacks both binding sites.

**Figure 6.**
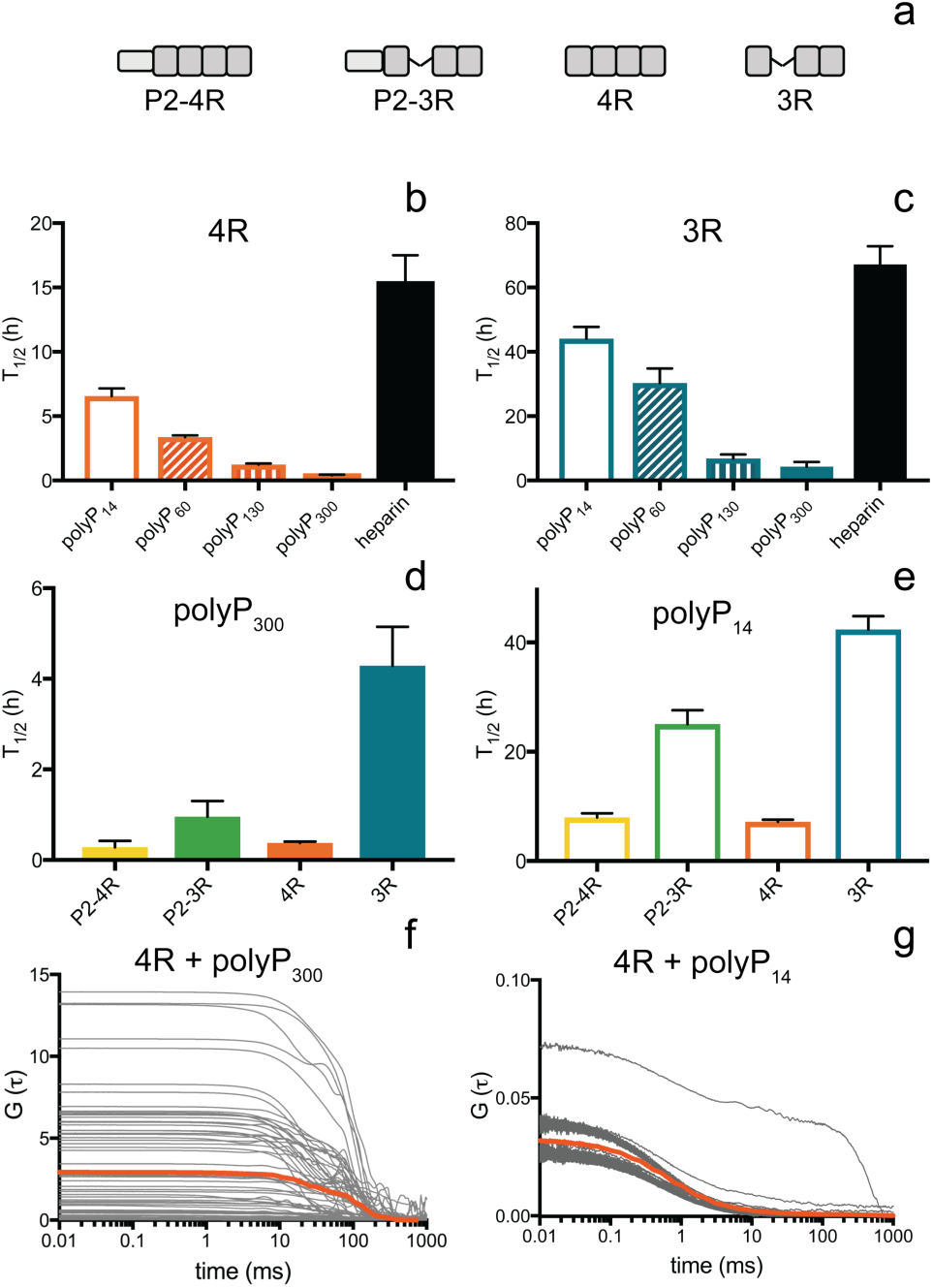
PolyP accelerates tau aggregation in a chain length dependent manner. The aggregation half-times (T_1/2_) of tau fragments (a) measured by ThT fluorescence comparing polyP chain lengths and heparin for 4R (b) and 3R (c) and comparing different fragments for polyP_300_ (d) and polyP_14_ (e). Autocorrelation curves of 4R in the presence of 1 mM polyP_300_ (f) or polyP_14_ (g). The individual autocorrelation curves are shown in gray and the average is in orange. Statistical analysis of repeat measurements for the ThT fluorescence assays is given in Table S5.

Our smFRET and aggregation data support the conclusion that binding of polyP to multiple intramolecular sites increases the efficacy of polyP in initiating tau aggregation. Moreover, our finding that longer polyP chains are more effective in stimulating tau aggregation suggest that longer polyP chains are able to interact with several tau monomers, thereby serving as an intermolecular scaffold to non-covalently cross-link tau monomers. FCS and smFRET measurements are typically carried out at pM to nM protein concentrations, which strongly disfavor intermolecular protein interactions. To directly test whether polyP is able to non-covalently cross-link tau monomers, we conducted FCS measurements at concentrations of 4R and polyP that are comparable to our ensemble aggregation experiments. The addition of 25 µM unlabeled 4R to 20 nM labeled 4R did not result in a change in its diffusion time, indicating that tau-tau interactions are not favored at the increased protein concentration in the absence of polyP (Fig. S7a and b). With the addition of 1 mM polyP_300_, however, extremely heterogeneous autocorrelation curves were recorded, suggesting the formation of large oligomeric assemblies or aggregates (Fig. 6f) (7). In contrast, the addition of 1 mM polyP_14_ resulted in only a few aberrant autocorrelation curves, reflecting fewer large species (Fig. 6g). These results suggest that longer polyP chains enhance intermolecular tau-tau interactions, which might play an important role in the acceleration of tau aggregation.

### PolyP is more effective at accelerating tau aggregation than heparin

Heparin is the most commonly used inducer of tau aggregation *in vitro* (5). To investigate whether heparin and polyP trigger similar changes in the conformational ensemble of tau, we compared their effects on tau in smFRET experiments. We measured two 2N4R constructs, tau_C17-C149_ and tau_C149-C372_, using concentrations of heparin (MW ∼17,000g) of 130 nM and 1.75 µM, which have the same equivalent charge as ∼5 µM and 80 µM polyP, respectively. At these concentrations, heparin induced shifts in the mean ET_eff_ in the same direction as polyP, although the magnitude of the shift was reproducibly smaller (Fig. 7; Table S6). Moreover, heparin does not appear to be as effective scaffold for binding multiple tau monomers (Fig. S7c), as polyP_300_ and heparin-induced aggregation of tau displayed a significantly slower kinetics than measured in the presence of polyP (Fig. 6b and 6c). Together, these data demonstrate that compared to heparin, polyP is significantly more effective at i) populating a more compact, aggregation-prone conformational ensemble of tau, ii) cross-linking multiple tau monomers, and consequently iii) inducing tau aggregation.

**Figure 7.**
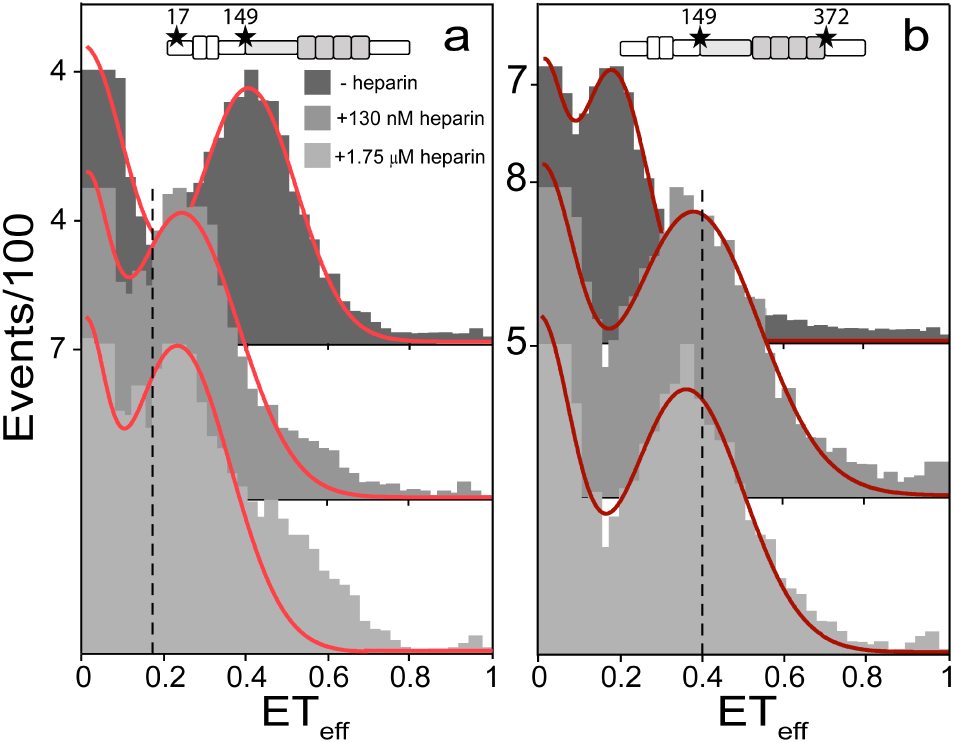
Heparin binding causes a smaller conformational change than polyP. Histograms from smFRET measurements probing two different regions of 2N4R, the N-terminal domain (C17-C149) (a) and the PRR and MTBR (C149-C372) (b) in the absence or presence of 130 nM or 1.75 µM heparin. The dotted black lines indicate the average ET_eff_ of the construct with 20 µM polyP_300_. At least three separate measurements of each condition and construct were made. The histograms shown are representative. Statistical analysis of repeat measurements is given in Table S6.

### PolyP competes with tubulin for tau binding

*In vitro*, polyP binds to tau and is very effective in accelerating its aggregation. In neurons, there are many other biomolecules that can compete with polyP for binding to tau. The most relevant of these is tubulin, tau’s primary cellular binding partner. We therefore decided to investigate whether polyP is a competitive binding partner of tau that is capable of affecting tau’s ability to interact with tubulin *in vitro*. For these experiments, we used the P2-4R fragment, which undergoes conformational changes (Fig. 4) and aggregates readily (Fig. 6) upon binding polyP and is known to interact with both soluble tubulin (34) and microtubules (35) *in vitro*. As a reporter for tubulin binding, we measured the diffusion time of fluorescently labeled P2-4R in the absence and presence of 5 µM tubulin by FCS (Fig. 8; Fig. S8). Consistent with our previous work, binding of P2-4R to tubulin resulted in an increase in the diffusion time from ∼0.66 ms to ∼0.93 ms (36). Upon titration with polyP_300_, we observed a polyP concentration-dependent decrease in the diffusion time, reflecting competitive displacement of tubulin from P2-4R by polyP (Fig. 8). In the presence of 100 µM polyP_300_, the diffusion time of P2-4R was comparable to that measured for P2-4R with saturating concentrations of polyP in the absence of tubulin (Fig. 3), suggesting that polyP had effectively replaced all of the bound tubulin.

**Figure 8.**
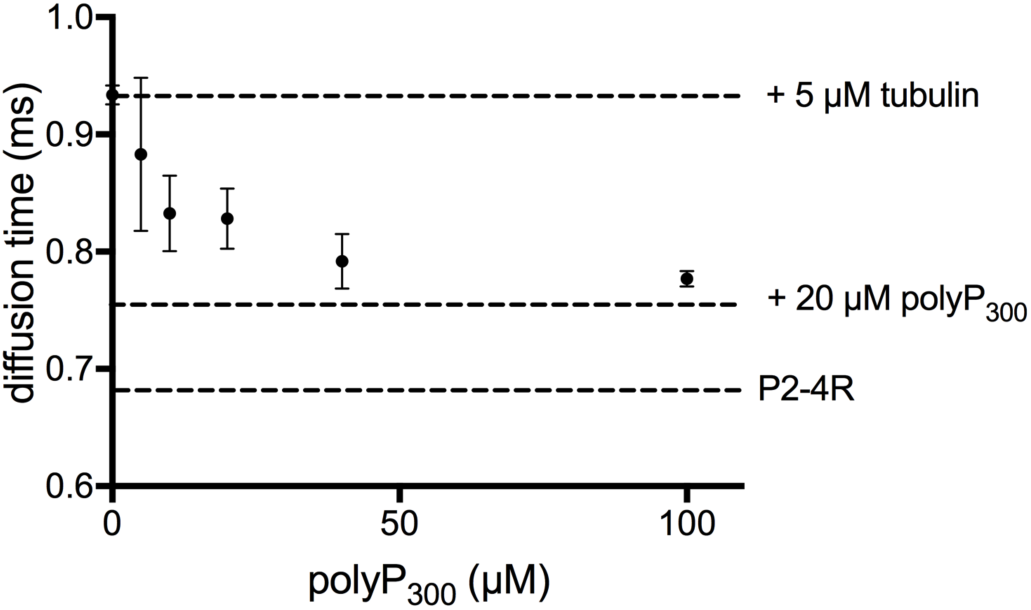
PolyP competes with tubulin for tau binding. FCS measurements of tubulin-bound P2-4R in the absence or presence of increasing concentrations of polyP_300_. For reference, the dashed lines correspond to the diffusion times measured for: P2-4R with 5 µM tubulin (upper), P2-4R with 20 µM polyP_300_ (middle) and P2-4R in buffer (lower). Data points are the mean and s.e.m of three independent measurements.

## Discussion

Our results identified three aspects of the interaction between tau and polyP relevant to its mechanism of enhancing tau aggregation: (1) binding of polyP to tau’s MTBR and PRR regions, which changes the conformational ensemble of monomer tau by causing compaction of those domains; (2) screening of electrostatic interactions between tau domains and disruption of long range interactions between tau’s termini and MTBR; and (3) facilitating intermolecular association of tau monomers (Fig. 9).

**Figure 9.**
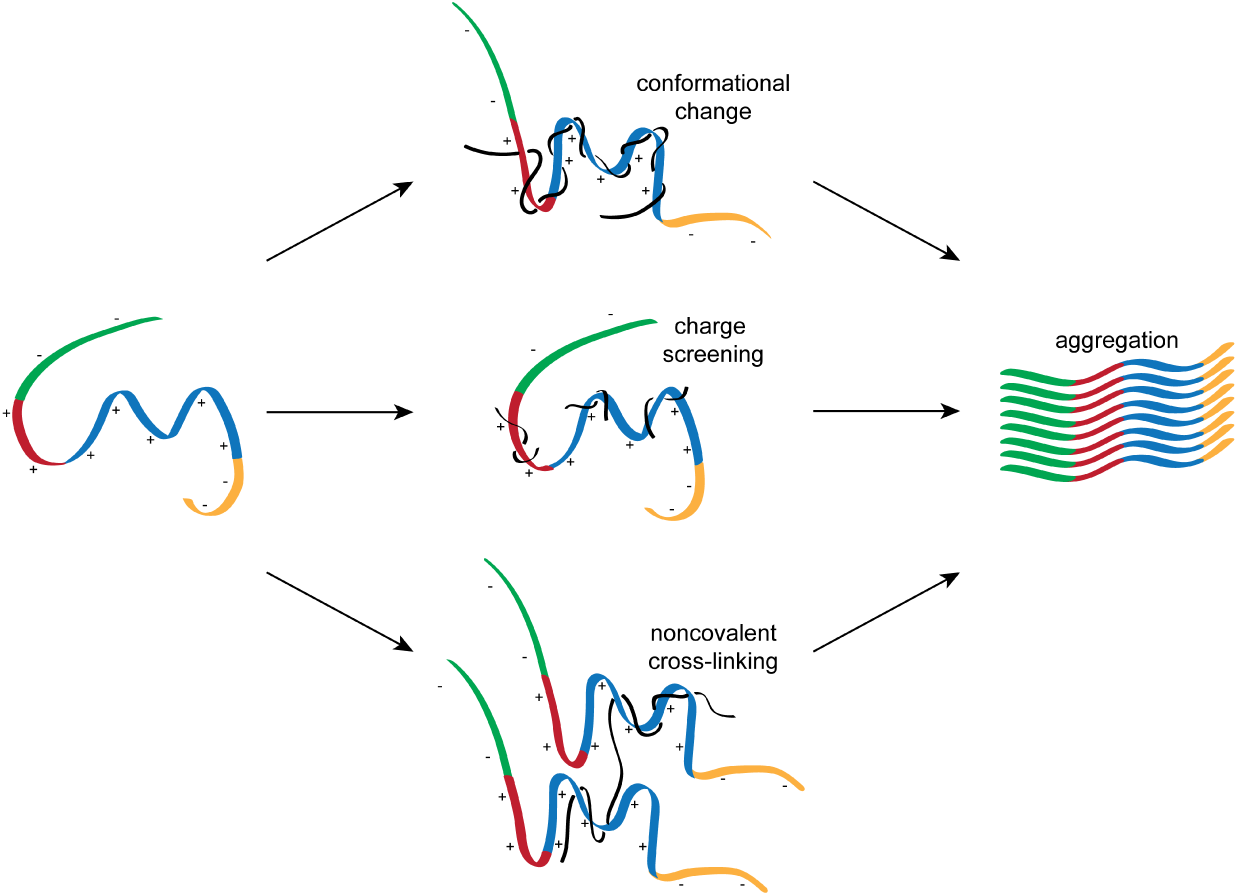
Proposed model of polyphosphate induced tau aggregation. The binding of polyP results in large conformational changes in monomer tau, charge screening and cross-linking between monomers. These combined effects result in the acceleration of tau aggregation. Tau domains are colored as: N-terminal domain (green), PRR (red), MTBR (blue) and C-terminal domain (yellow). PolyP is shown in black.

SmFRET measurements of full-length tau constructs most clearly illustrated that polyP binding causes both local (compaction of the PRR and MTBR) and long-range (loss of interaction between the termini and central region) changes in tau’s conformational ensembles (Fig. 2). We found that polyP binds to both PRR and MTBR regions, yet compaction of the MTBR domain is only observed in the presence of the PRR (compare Fig. 4b and c). We reason that the observed changes in the conformational ensemble result from the noncovalent intramolecular cross-linking of binding sites in the PRR and MTBR by a single polyP chain (Fig. 9). Further evidence in support of this idea came from the use of polyP of different chain lengths. Even at saturating concentrations, we found that polyP_14_, which is too short to span between binding sites in the PRR and MTBR (37), causes only a slight shift in the mean ET_eff_ of full-length tau (Fig. 5) while longer chain lengths (polyP_60_, polyP_130_ and polyP_300_) cause much larger shifts. Interestingly, the shift caused by polyP_60_ is to a higher mean ET_eff_ value than that of polyP_130_ and polyP_300_ for tau_C149-C372_ (Fig. 5b). The shorter end-to-end distance of polyP_60_ may not be able to span between binding sites without imposing a more compact conformational ensemble on tau. This effect is likely only seen for the tau_C149-C372_ construct (Fig. 5b) but not tau_C17-C149_ (Fig. 5a) because only the probes for the former encompass the PRR and MTBR binding sites.

We consider that both charge screening and intermolecular cross-linking are likely relevant to the general capability of polyP to accelerate tau aggregation. The tau fragments are highly positively charged, disfavoring intermolecular interactions in solution. Indeed, FCS measurements with high concentrations of unlabeled monomer 4R show no evidence of tau assembly (Fig. S7). Binding of negatively charged polyP decreases the electrostatic repulsion between monomers so that intermolecular interactions become more favorable and binding of polyP to tau is effectively blocked by increasing the buffer ionic strength (Fig. S3). While all polyP chain lengths are capable of accelerating aggregation through this mechanism (Fig. 6b), longer polyPs are more effective because they are also capable of facilitating binding to multiple tau monomers, effectively increasing their local concentration (Fig. 6f, g and Fig. 9). The fact that polyP most effectively accelerates the aggregation of tau fragments that contain both P2, in the PRR, and R2, in the MTBR suggests that the presence of additional binding sites may facilitate intermolecular scaffolding by polyP.

One challenge faced by in vitro aggregation studies is translating their results to a more physiological context. For example, while disease models generally describe alterations to tau (such as mutations found in tauopathies) as enhancing the aggregation propensity of tau, the effects *in vitro* are usually fairly moderate (1, 38). Natively, tau is associated with either microtubules (35, 39) or soluble tubulin (40), although this interaction is likely to be highly dynamic (25, 26). We find that polyP is able to displace tubulin from tau (Fig. 8) at concentrations much lower than the reported cytosolic concentrations of polyP (41). The cellular cytoplasm is much more complex than the tertiary mixture explored here and thus whether polyP could compete for binding to tau under physiological conditions remains to be tested. However, our measurements strongly suggest that polyP is capable of binding tau in the presence of tubulin or microtubules. Moreover, under pathological conditions where tau binding to microtubules is compromised by mutation or hyperphosphorylation (42), the cytoplasmic pool of tau available to interact with polyP is increased. Future studies will investigate whether tubulin-dissociated, polyP-bound monomer might be putative target for therapeutics (reviewed in (43). Our results here provide insight into the conformational features of this monomer and may eventually aid in the design of therapeutics to combat tauopathies.

## Author Contributions

S.P.W, J.L., U.J. and E.R. designed the experiments; S.P.W., H.E.M., J.L. and J.M performed research and analyzed data; S.P.W., U.J. and E.R. wrote the manuscript.

## Acknowledgements

We thank K.M. McKibben and H.Y.J. Fung for providing some of the single-labeled tau and unlabeled constructs. This work was funded by the National Institute of Health NIHAG053951 (to E.R), NR35GM122506 and R21AG055090 (to U.J), PhD fellowship of the Boehringer Ingelheim Fonds, Stiftung fuer medizinische Grundlagenforschung (to J.L) and The Vagelos Program in the Molecular Life Sciences at the University of Pennsylvania (to H.M).

